# Extensive protected area coverage and an updated global population estimate for the Endangered Madagascar Serpent-eagle identified from species-habitat associations using remote sensing data

**DOI:** 10.1101/2022.04.26.489510

**Authors:** Luke J. Sutton, Armand Benjara, Lily-Arison Rene de Roland, Russell Thorstrom, Christopher J.W. McClure

## Abstract

Knowledge gaps regarding distribution, habitat associations, and population size for rare and threatened range-restricted taxa leads to uncertainty in directing conservation action. Quantifying range metrics and species-habitat associations using Species Distribution Models (SDMs) with remote sensing habitat data can overcome these setbacks by establishing baseline estimates for biological parameters critical for conservation assessments. Area of habitat (AOH) is a new range metric developed by the International Union for the Conservation of Nature (IUCN) Red List. AOH seeks to quantify inferred habitat within a species’ range to inform extinction risk assessments. Here, we use SDMs correlating occurrences with remote-sensing covariates, to calculate a first estimate of AOH for the Endangered Madagascar Serpent-eagle (*Eutriorchis astur*), and then update additional IUCN range metrics and the current global population estimate. From these baselines we then conduct a gap analysis assessing protected area coverage. Our continuous SDM had robust predictive performance (Continuous Boyce Index = 0.835) and when reclassified to a binary model estimated an AOH = 30,121 km^2^, 13 % less than the current IUCN range map. We estimate a global population of 533 mature individuals derived from the Madagascar Serpent-eagle AOH metric, which was within the current IUCN population estimates. The current protected area network covered 95 % of AOH, with the binary model identifying three key habitat areas as new protected area designations to fully protect Madagascar Serpent-eagle habitat. Our results demonstrate that correlating presence-only occurrences with remote sensing habitat covariates can fill knowledge gaps useful for informing conservation action. Applying this spatial information to conservation planning would ensure almost full protected area coverage for this endangered raptor. For tropical forest habitat specialists, we recommend that potential predictors derived from remote sensing, such as vegetation indices and biophysical measures are considered as covariates, along with other variables including climate and topography.

## Introduction

Mapping geographic ranges and identifying environmental requirements of threatened species are fundamental research areas in conservation biology (Riddle *et al*. 2011). Defining species’ spatial and ecological range limits is essential to assess the various threats facing many taxa in rapidly changing environments (Ladle & Whittaker 2011), and to formulate viable conservation plans for species survival (Margules & Pressey 2000; Sutton *et al*. 2021a). However, significant knowledge gaps still exist regarding the full area of distribution and environmental attributes of where individual species occur, commonly termed the ‘Wallacean Shortfall’ (Lomolino 2004). The Wallacean Shortfall contributes to a second knowledge deficit where, if the current range of a species is unknown or not fully described, it is not possible to determine whether and when a species is in decline or possibly gone extinct. Thus, the environmental factors that limit the distribution and abundance of many threatened species are still poorly understood (Marcer *et al*. 2013).

The current paradigm is that climate plays a central role in determining species distributions at broad scales (Pearson & Dawson 2003). However, recent work has demonstrated that biotic interactions (Aragón *et al*. 2018), landcover (Tuanmu & Jetz 2014, 2015), and topography (Meineri & Hylander 2017) are also important at setting range limits for many taxa. Species Distribution Models (SDMs) are a group of geospatial statistical methods that assess species-habitat associations and predict distribution based on correlating environmental covariates with species occurrences (Matthiopoulos *et al*. 2020). SDMs can be effective for estimating potential range limits and ecological associations using satellite remote sensing data coupled with occurrences from unstructured surveys and community science projects (Bradter *et al*. 2017). This includes for threatened species distributed across remote, hard to survey areas (Sutton *et al*. 2021b).

The endemic Madagascar Serpent-eagle (*Eutriorchis astur*) is a cryptic, medium-sized raptor with a restricted distribution across tropical forests in eastern Madagascar (BirdLife International 2016). The species is one of the rarest raptors globally and is currently classified as ‘Endangered’ on the International Union for Conservation of Nature (IUCN) Red List (BirdLife International 2016). This forest-dependent raptor was once considered extinct but was rediscovered three decades ago by Peregrine Fund biologists (Thorstrom *et al*. 1995). Madagascar Serpent-eagles generally prefer uninterrupted expanses of lowland and mid-altitude tropical forest, with habitat loss and fragmentation the primary threats to the species future persistence (Thorstrom & Rene de Roland 2000). Despite being termed a serpent eagle, snakes comprise only a small proportion of Madagascar Serpent-eagle prey, with chameleons and geckos accounting for > 80 % of diet (Thorstrom & de Roland 2000). Recent research suggests the Madagascar Serpent-eagle may be vulnerable to both climate change (Andriamasimanana & Cameron 2013) and increasing forest fragmentation (Benjara *et al*. 2021).

From surveys using playback techniques (Thorstrom & Rene de Roland 2000), the known range of the Madagascar Serpent-eagle is now thought to be considerably larger than previously estimated and it may not be as rare as once thought (BirdLife International 2016). However, the environmental determinants of Madagascar Serpent-eagle distribution and abundance are still largely unknown. The global population is still very small, estimated between 250-999 mature individuals, and is likely to be decreasing (BirdLife International 2016). Spatial modelling can therefore help determine the essential ecological requirements of the Madagascar Serpent-eagle and update range metrics and population size estimates, both currently identified as priority areas of research (Thorstrom *et al*. 1995; Thorstrom & Rene de Roland 2000; BirdLife International 2016). Further, predicting the distributional potential for the Madagascar Serpent-eagle would enable specific hypotheses to be developed and tested on the processes limiting its distribution. This includes directing current field sampling protocols to identify potential areas of occupation (*sensu* Peterson & Anamza 2015).

Improving the predictive power of spatial models by incorporating biotic, landcover, and topographical predictors derived from satellite remote sensing would also lead to higher certainty on where to designate new protected areas and strengthen the existing protected area network (Elith & Leathwick 2009). Applying this knowledge to current conservation management can then direct designation of protected areas in line with suitable environmental areas (Sutton *et al*. 2022). Given this background, our aims are to apply spatial predictive modelling to estimate distribution and identify ecological range limits for the Madagascar Serpent-eagle. Our key objective is to use this information to inform current spatial conservation planning and estimate a potential population size. Here, we set out baseline estimates for: (**1**) the current distribution of the Madagascar Serpent-eagle based on remote sensing habitat covariates, (**2**) identify range-wide species-habitat associations, and (**3**) update IUCN range metrics and a population size estimate. Last, from these baselines we then calculate protected area coverage, and conduct a gap analysis to identify priority designations for new protected areas.

## Methods

### Study extent and species locations

We defined the species’ accessible area (Barve *et al*. 2011), as the ecoregions corresponding to both Madagascar lowland and subhumid tropical forest extracted from the World Wildlife Fund (WWF) terrestrial ecoregions shapefile (Fig. 1; Olson *et al*. 2001). We masked out the remaining ecoregions in the far north, east, and south of Madagascar, because the Madagascar Serpent-eagle is a habitat specialist of moist tropical forest (Thorstrom & Rene de Roland 2000; Benjara *et al*. 2021) and has not been recorded outside of these ecoregions. We compiled a database of 33 Madagascar Serpent-eagle point localities from the Global Raptor Impact Network (GRIN), a global population monitoring information system for all raptors (McClure *et al*. 2021). For the Madagascar Serpent-eagle, GRIN consists of locations from unstructured surveys which only recorded presence (*n* = 24), a literature search (*n* = 4, Sheldon & Duckworth 1990; Raxwothy & Colston 1992; Hawkins *et al*. 1998; Karpanty & Grella 2001) and community science data from the Global Biodiversity Information Facility (*n =* 5; GBIF 2020) (See Supplementary Material).

**Figure 1.**
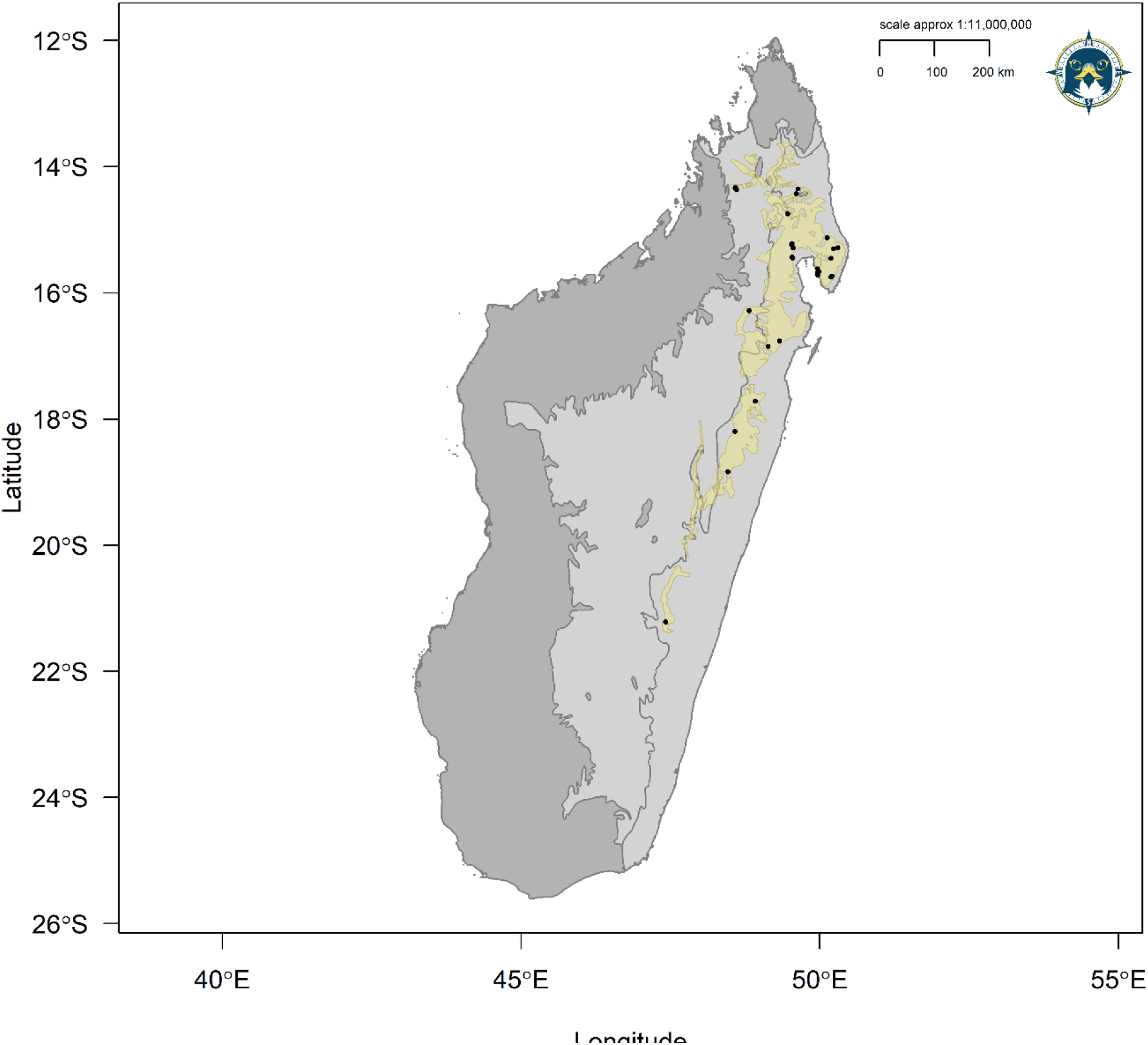
Range map for the Madagascar Serpent-eagle with our model accessible area from the tropical moist forest ecoregions (light grey) and IUCN range map (khaki). Black points define Madagascar Serpent-eagle occurrences from the Global Raptor Impact Network population monitoring system.

### Habitat covariate models

We considered eight potential habitat covariates *a prioiri* related empirically to known Madagascar Serpent-eagle habitat associations (Thorstrom & Rene de Roland 2000; Benjara *et al*. 2021). These were derived from satellite remote sensing products representing climate, landcover, topography, and vegetation at a spatial resolution of 30 arc-seconds (∼1 km resolution; Table 1). We downloaded raster layers from the EarthEnv (www.earthenv.org), ENVIREM (Title & Bemmels 2018), and Dynamic Habitat Indices (Radeloff 2019) repositories, which were then cropped and masked to a delimited polygon representing the species accessible area (Fig. 1). Climatic Moisture Index is a scaled measure (−1 ≤ Climatic Moisture Index ≤ 1) of the ratio of annual precipitation and annual evapotranspiration (Willmott & Feddema 1992) used here as a proxy for moist tropical forest coverage. Evergreen Forest is a measure of percentage landcover here representing broadleaf tropical evergreen forest derived from consensus products integrating GlobCover (v2.2), MODIS land-cover product (v051), GLC2000 (v1.1), and DISCover (v2) from the years 1992-2006. Full details on methodology and image processing for evergreen forest can be found in Tuanmu & Jetz (2014). 1452

**Table 1.** Habitat covariates selected *a priori* and considered as potential covariates used in all spatial analyses for the Madagascar Serpent-eagle. FPAR = Fraction of absorbed Photosynthetically Active Radiation, NDVI = Normalized Difference Vegetation Index.

| Covariate | Source | Citation | Year(s) |
| --- | --- | --- | --- |
| Climatic Moisture Index | ENVIREM | Title & Bemmels 2018 | 2000 |
| Elevation | EarthEnv | Amatulli *et al.* 2018 | 2010 |
| Evergreen forest | EarthEnv | Tuanmu & Jetz 2014 | 1992-2005 |
| FPAR | DHI | Radeloff *et al.* 2019 | 2003-2014 |
| Heterogeneity | EarthEnv | Tuanmu & Jetz 2015 | 2001-2005 |
| Leaf Area Index | DHI | Radeloff *et al.* 2019 | 2003-2014 |
| NDVI | DHI | Radeloff *et al.* 2019 | 2003-2014 |
| Terrain Roughness Index | ENVIREM | Title & Bemmels 2018 | 2000 |

Heterogeneity is a biophysical texture measure closely related to vegetation structure, composition, and diversity (i.e., species richness) derived from textural features of Enhanced Vegetation Index between adjacent pixels; sourced from the Moderate Resolution Imaging Spectroradiometer (MODIS, https://modis.gsfc.nasa.gov/). We inverted the raster cell values in the original EarthEnv variable ‘Homogeneity’ (Tuanmu & Jetz 2015) to represent the spatial variability and arrangement of vegetation species richness on a continuous scale which varies between zero (minimum heterogeneity, low species richness) and one (maximum heterogeneity, high species richness). Elevation was derived from a digital elevation model product from the 250m Global Multi-Resolution Terrain Elevation Data 2010 (Danielson & Gesch 2011). Terrain Roughness Index is a measure of variation in topography around a central pixel, with lower values indicating flat terrain and higher values indicating larger differences in elevation of neighbouring pixels (Wilson *et al*. 2007).

Last, we used three biophysical vegetation layers based on averaged 8 and 16-day MODIS vegetation products, used here as composite Dynamic Habitat Index (DHI) products (Radeloff 2019). We used the single composite phenology curve product for each DHI vegetation layer, summarising three measures of vegetation productivity between 2003-2014: annual cumulative, minimum throughout the year, and seasonality as the annual coefficient of variation. Normalized Difference Vegetation Index (NDVI) provides a measure of photosynthetic activity, linked to species richness and productivity (Huete *et al*. 2002). However, NDVI can saturate in dense vegetation and highly productive areas (such as moist tropical forests) and cannot distinguish differences in productivity in these areas (Huete *et al*. 2002). Therefore, we used two further measures that directly assess productivity, providing a more accurate proxy for vegetation coverage: Leaf Area Index and Fraction of absorbed Photosynthetically Active Radiation (FPAR).

Both Leaf Area Index and FPAR incorporate landcover in their calculation and use reflectance values from up to seven MODIS bands, compared to the two or three bands for NDVI and Enhanced Vegetation Index respectively (Hobi *et al*. 2017). Leaf Area Index is a measure of the amount of foliage within the plant canopy and a key driver of primary productivity (Asner *et al*. 2003). FPAR is a measure of productivity inferred from available photosynthetic activity driven by solar radiation (Myneni *et al*. 2002), characterising the energy absorption of the vegetation canopy. Leaf Area Index and FPAR are closely related measures, with Leaf Area Index recommended for high productivity areas and FPAR for lower productivity areas (Radeloff 2019). Combined, we used each Dynamic Habitat Index as a proxy for food availability, assuming that summarising vegetation productivity annually over the 11-year period captures seasonal variations in species richness that Madagascar Serpent-eagles would use as food (Hobi *et al*. 2017).

We selected covariates to use in our final model based on an information theoretic approach using Akaike’s Information Criterion (Akaike 1974) corrected for small sample sizes (AIC_c_; Hurvich & Tsai 1989) in the R package AICcmodavg (Mazerolle 2020). We fitted six candidate models using logistic regression with a binomial error term and logit link function using generalised linear models (GLMs) in the R package stats (R Core Team, 2018). All our candidate models were fitted to derive maximum likelihood estimates on model parameters significantly different from zero, with no interaction terms. Predictors were standardized with a mean of zero and standard deviation of one. Because the occurrence data correspond to a presence-only dataset, background availability was randomly sampled using 10,000 pseudo-absence points suitable for regression-based modelling (Barbet-Massin *et al*. 2012). We assigned equal weights to both presence and background points allowing consistent sampling across the model calibration area. We did this to avoid saturating the models with excessive absence weighting, which makes presence trends difficult to detect (Elith & Leathwick 2007).

First, we fitted model 1 with all eight covariates representing climate, landcover, topography, and vegetation, model 2 with only landcover, topographic, and vegetation variables, and model 3 with landcover and vegetation plus elevation, but without Terrain Roughness Index. We fitted model 4 only considering landcover and vegetation variables, and finally models 5 and 6 fitted with landcover and vegetation both with and without NDVI and FPAR. We did not include an intercept-only model because its ΔAICc score was not competitive (ΔAIC_c_ = 20.75). We fitted linear terms to all model covariates except for Climatic Moisture Index and Terrain Roughness Index which were fitted with quadratic terms because we expected a quadratic effect with values of both covariates to be highest at intermediate values and decrease at lower and higher values. All candidate models with a ΔAIC_c_ < 2 were considered as having strong support (Burnham & Anderson 2004), and the best supported model was selected using the lowest ΔAIC_c_ and highest AIC_c_ weighting. Last, we tested the covariates for multicollinearity directly at the Madagascar Serpent-eagle occurrences from our best supported model considering Variance Inflation Factors <2 (Dormann *et al*. 2013).

### Species Distribution Models

After identifying the most parsimonious model covariates using binomial GLMs, we fitted candidate SDMs, further tuning model parameters using penalized elastic net logistic regression in the R package maxnet (Phillips *et al*. 2017). Penalizing model coefficients reduces model variance, resulting in a regression model that generalizes better than standard logistic regression (Valavi *et al*. 2021). Penalized logistic regression imposes a regularization penalty to the model coefficients reducing model complexity by shrinking the covariates that contribute the least to model prediction (Gastón & García-Viñas 2011; Fithian & Hastie 2013). An elastic net is used to perform automatic variable selection and continuous shrinkage simultaneously (via the *glmnet* package, Friedman et al. 2010), retaining all covariates that contribute less by shrinking coefficients to either exactly zero or close to zero. We fitted SDMs via maximum penalized likelihood estimation using a complementary log-log (cloglog) link function as a continuous index of environmental suitability, with 0 = low suitability and 1 = high suitability. We parametrized the penalized logistic regression model using infinite weighting (presence weights = 1, background = 100) equivalent to an inhomogeneous Poisson process because this is the most effective method to model presence-background data as used here (Warton & Shepherd 2010).

We used a random sample of 10,000 background points as pseudo-absences recommended for regression-based modelling (Barbet-Massin *et al*. 2012) and to sufficiently sample the background calibration environment (Guevara *et al*. 2018). Optimal-model selection was based on Akaike’s Information Criterion (Akaike 1974) corrected for small sample sizes (AIC_c_; Hurvich & Tsai 1989), to determine the most parsimonious model from two model parameters: regularization beta multiplier (β; level of coefficient penalty) and feature classes (response functions; Warren & Seifert 2011). Twenty-seven candidate models of varying complexity were built by conducting a grid search using a range of regularization multipliers from 1 to 5 in 0.5 increments, and three feature classes (response functions: Linear, Quadratic, Hinge) in all possible combinations using the ‘jackknife’ method of *k-*fold cross validation within the R package ENMeval (Muscarella *et al*. 2014).

The *n* – 1 jackknife cross validation approach is specifically used to test predictions using small occurrence datasets where the number of *k* folds is equal to the number of occurrences (*n*). All records are used but one in each model iteration, rather than losing valuable records via data splitting, with the single withheld record used once for testing (Gerstner *et al*. 2018). From each withheld test record, *n* models are calibrated and evaluated at each iteration across all *n* models (Shcheglovitova & Anderson 2013). All models with a ΔAIC_c_ < 2 were considered as having strong support (Burnham & Anderson 2004), with the model that had the lowest ΔAIC_c_ that used all three feature classes selected as the best supported model. Response functions and parameter estimates were used to measure variable performance within the optimal calibration SDM. We used Continuous Boyce Index (Hirzel *et al*. 2006) as a measure of how predictions differ from a random distribution of observed presences (Boyce *et al*. 2002). Last, we tested the optimal model against random expectations using partial Receiver Operating Characteristic ratios (pROC), which estimate model performance by giving precedence to omission errors over commission errors (Peterson *et al*. 2008) (See Supplementary Material).

### Range metrics and population size estimation

We followed the spatial framework of Sutton *et al*. (2022) and converted the final range-wide continuous prediction into a binary threshold prediction which we term *model* area of habitat (AOH), so as to be distinct from the standard IUCN AOH methodology (Brooks *et al*. 2019). To calculate *model* AOH in suitable pixels we reclassified the continuous prediction to a binary threshold using all pixel values equal to or greater than maximizing the sum of sensitivity and specificity (maxTSS) from the continuous model prediction. We used maxTSS because it is the most appropriate threshold for SDM conservation applications using presence-only data (Liu *et al*. 2013). We calculated two further IUCN range metrics from our *model* AOH binary prediction in the R package redlistr (Lee *et al*. 2019). To do this we first converted the *model* AOH raster to a polygon using an 8-neighbour patch rule and applied a smoothing function using the Chaikin algorithm (Chaikin 1974) in the R package smoothr (Strimas-Mackey 2021).

First, we calculated Extent of Occurrence (EOO), fitting a minimum convex polygon around the furthest boundaries of the smoothed *model* AOH polygon following IUCN guidelines (IUCN 2018). We calculated both a maximum EOO, including all the area with the minimum convex polygon, and a minimum EOO, masking out the areas that could either not be occupied, or are unlikely to be, within the minimum convex polygon, in our case over the ocean and outside of the moist tropical forest ecoregions (Marcer *et al*. 2013). Second, we calculated Area of Occupancy (AOO) as the number of raster pixels predicted to be occupied, scaled to a 2×2 km grid (4-km^2^ cells) following IUCN guidelines (IUCN 2018). All range metric calculations were performed using a Transverse cylindrical equal area projection following IUCN guidelines (IUCN 2018).

Finally, we calculated the number of Madagascar Serpent-eagle pairs our *model* AOH could support as directly proportional to the available habitat required by a territorial pair. We defined the habitat area for a breeding pair based on nearest neighbour distances of 6-km between nests from the Masoala Peninsula, which currently has the highest known density of breeding Madagascar Serpent Eagles (Thorstrom & Rene de Roland 2000). We used the area of a circle (113 km^2^) calculated from the 6-km inter-nest distance and then divided our *model* AOH area by this breeding habitat area to estimate the total number of mature individuals across the species range using the IUCN Red List definitions for population size (IUCN 2019). Finally, we divided that figure by 2 to give the number of potential breeding pairs.

### Protected area coverage

We assessed the level of protected area coverage using the World Database of Protected Area terrestrial shapefile for Madagascar (as of December 2021; UNEP-WCMC & IUCN 2021). We quantified how much protected area representation is needed for the Madagascar Serpent-eagle dependent on the *model* AOH to calculate a protected area ‘representation target’ following the formulation of Rodrigues *et al*. (2004),

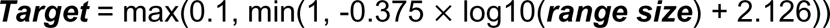

where ‘***Target****’* is equal to the percentage of protected target representation required for the species ‘***range size’***. We calculated the difference between the current level of protected area coverage compared to the target level representation using the *model* AOH intersected with the protected area polygons establishing those protected areas covering areas of habitat suitability ≥ maxTSS threshold. The protected area network polygons were then overlaid with the binary map identifying gaps in habitat suitability ≥ maxTSS threshold which were not covered by the terrestrial protected area polygons. Model development and geospatial analysis were performed in R (v3.5.1; R Core Team, 2018) using the raster (Hijmans 2017), rgdal (Bivand *et al*. 2019), rgeos (Bivand & Rundle 2019) and sp (Bivand *et al*. 2013) packages.

## Results

### Habitat covariate models

Three candidate GLMs had strong support with an ΔAIC_c_ < 2 (Table 2), with our best supported candidate GLM, model 6 (Heterogeneity + Evergreen Forest + Leaf Area Index + NDVI), with half as much AICc weighting (AIC_c_ *w =* 0.44) from the next best supported candidate GLM, model 5 (AIC_c_ *w =* 0.29). From the best supported GLM linear beta coefficients (Table S1; Fig. 2), NDVI had the strongest positive association with Serpent-eagle occurrence (β = 2.128, ns), followed by Evergreen Forest (β = 1.802, *p* <0.01) and Heterogeneity (β = 1.004, ns). Leaf Area Index had the strongest negative association with Serpent-eagle occurrence (Fig. 2). The covariates from the best supported GLM model all had low collinearity (VIF <2; Table S2, Fig. S1) and thus all covariates were included in the penalized SDMs. 2104

**Figure 2.**
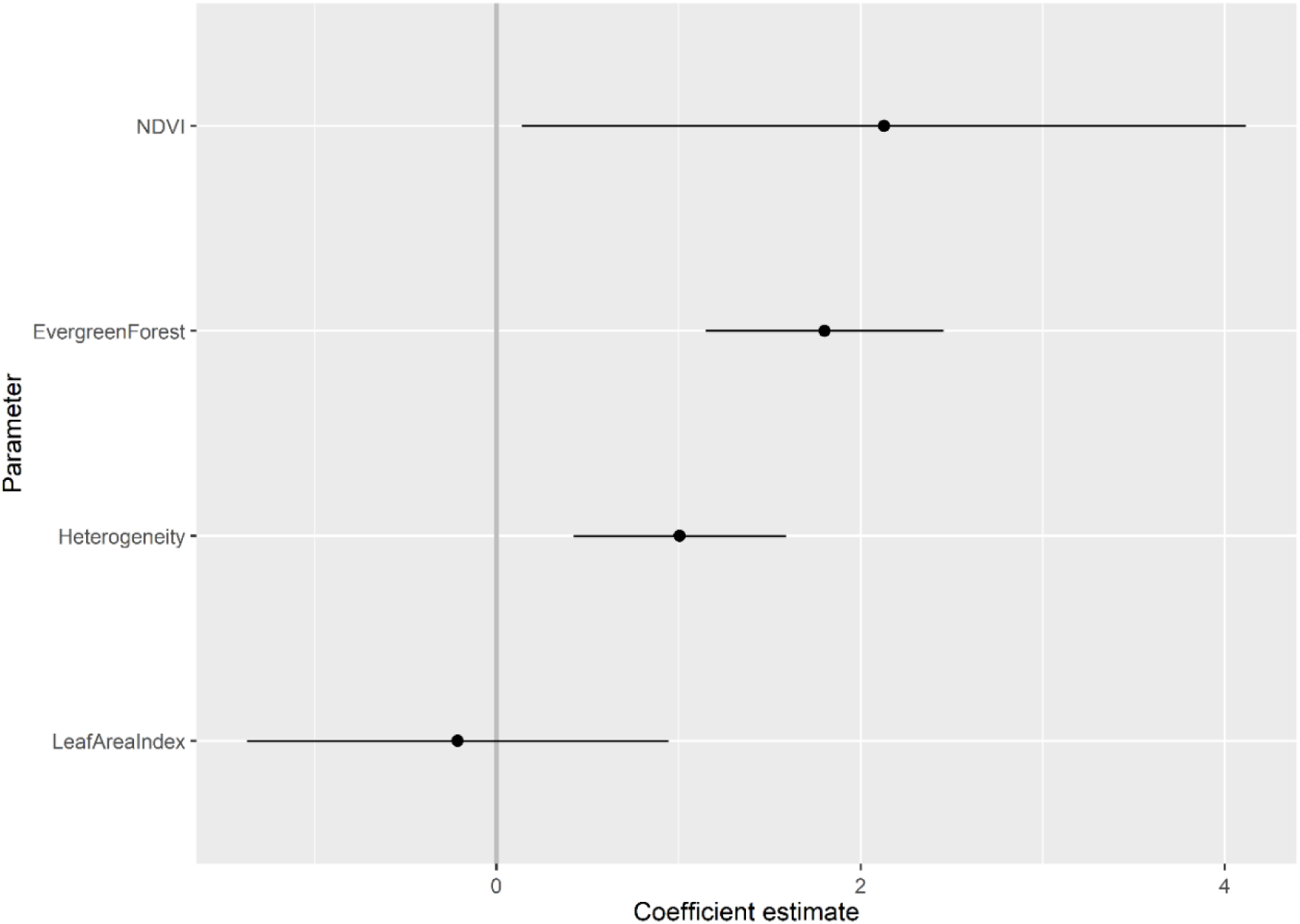
Coefficient estimates (with standard errors) for the best supported candidate logistic regression habitat covariate model (#6) for the Madagascar Serpent-eagle from the most parsimonious model using Akaike’s Information Criterion corrected for small sample sizes.

**Table 2.** Comparison of candidate logistic regression habitat covariate models for the Madagascar Serpent-eagle using Akaike’s Information Criterion corrected for small sample sizes (AICc). Number of model parameters (*K*), change in AICc (ΔAICc), Akaike weight (AICc *w*) and log-likelihhod (LL) are reported for each candidate model. CMI = Climatic Moisture Index; TRI = Terrain Roughness Index; HG = Heterogeneity; EF = Evergreen Forest; ELEV = Elevation; LAI = Leaf Area Index; NDVI = Normalized Difference Vegetation Index; FPAR = Fraction of absorbed Photosynthetically Active Radiation.

| # | Candidate model | $K$ | $AIC_c$ | $\Delta AIC_c$ | $AIC_c w$ | LL |
| --- | --- | --- | --- | --- | --- | --- |
| 6 | HG + EF + LAI + NDVI | 5 | 24.22 | 0.00 | 0.44 | -7.11 |
| 5 | HG + EF + LAI + FPAR | 5 | 25.05 | 0.82 | 0.29 | -7.52 |
| 4 | HG + EF + LAI + NDVI + FPAR | 6 | 26.22 | 2.00 | 0.16 | -7.10 |
| 3 | HG + EF + ELEV + LAI + NDVI + FPAR | 7 | 28.20 | 3.97 | 0.06 | -7.09 |
| 2 | $TRI^2$ + HG + EF + ELEV + LAI + NDVI + FPAR | 8 | 29.84 | 5.62 | 0.03 | -6.91 |
| 1 | $CMI^2$ + $TRI^2$ + HG + EF + ELEV + LAI + NDVI + FPAR | 9 | 31.67 | 7.45 | 0.01 | -6.83 |

### Species Distribution Models

Three candidate SDMs had a ΔAIC_c_ ≤ 2, with the best supported penalized SDM using linear, quadratic and hinge terms and a coefficient penalty β = 3 as model parameters (model 15, Table S3). The optimal SDM had good calibration accuracy (CBI = 0.835) and was robust against random expectations (pROC = 1.892, SD± 0.058, range: 1.746 – 2.000). The largest continuous area of habitat extended along the remaining areas of tropical moist forest of the Eastern Malagasy Region in the Central and Eastern domains (Fig. 3). A second substantial area of habitat was identified across the Masoala Peninsula and further north into forested, lower elevation areas of the Tsaratanana Massif. 113

**Figure 3.**
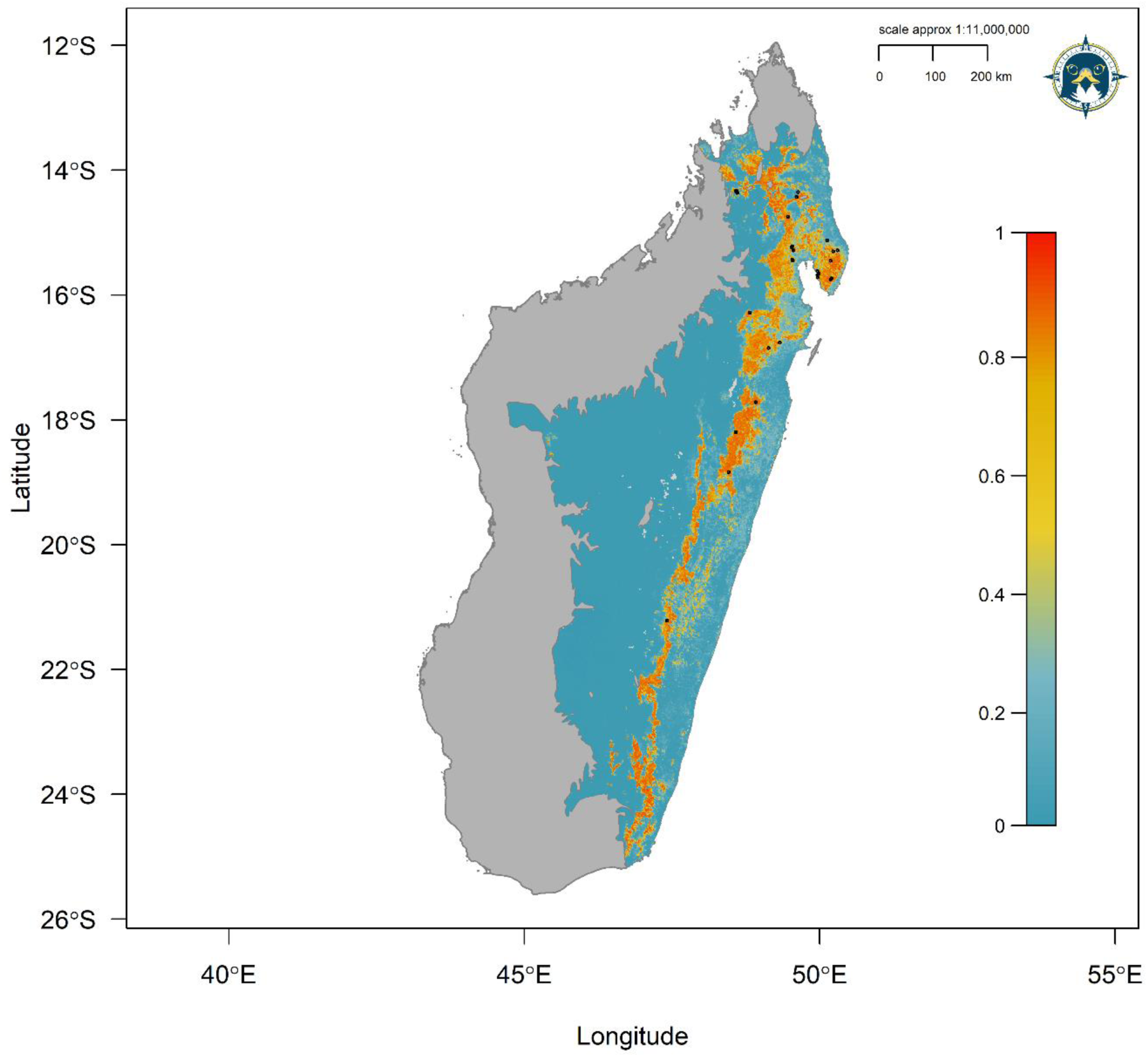
Continuous Species Distribution Model for the Madagascar Serpent-eagle using a penalized logistic regression model algorithm. Map denotes continuous prediction with red areas (values closer to 1) having highest habitat suitability, orange/yellow medium suitability and blue/green low suitability. Black points define Madagascar Serpent-eagle occurrences from the Global Raptor Impact Network population monitoring system.

The optimal model shrinkage penalty was able to retain four non-zero beta coefficients, setting to zero most model terms, meaning only a small subset of covariate terms were highly informative to model prediction (Figs. S2-S4). From the penalized linear beta coefficients, the Madagascar Serpent-eagle was most positively associated with vegetation heterogeneity (1.220), followed by NDVI (0.148), Evergreen Forest (0.043) and Leaf Area Index (0.002). From the penalized response functions, peak suitability for vegetation heterogeneity was at 90-95 %, with highest suitability for composite NDVI values > 20 (Fig. 4). The Madagascar Serpent-eagle was positively associated with > 95 % Evergreen Forest cover with a flat response to Leaf Area Index values between 0.0-3.0. 113

**Figure 4.**
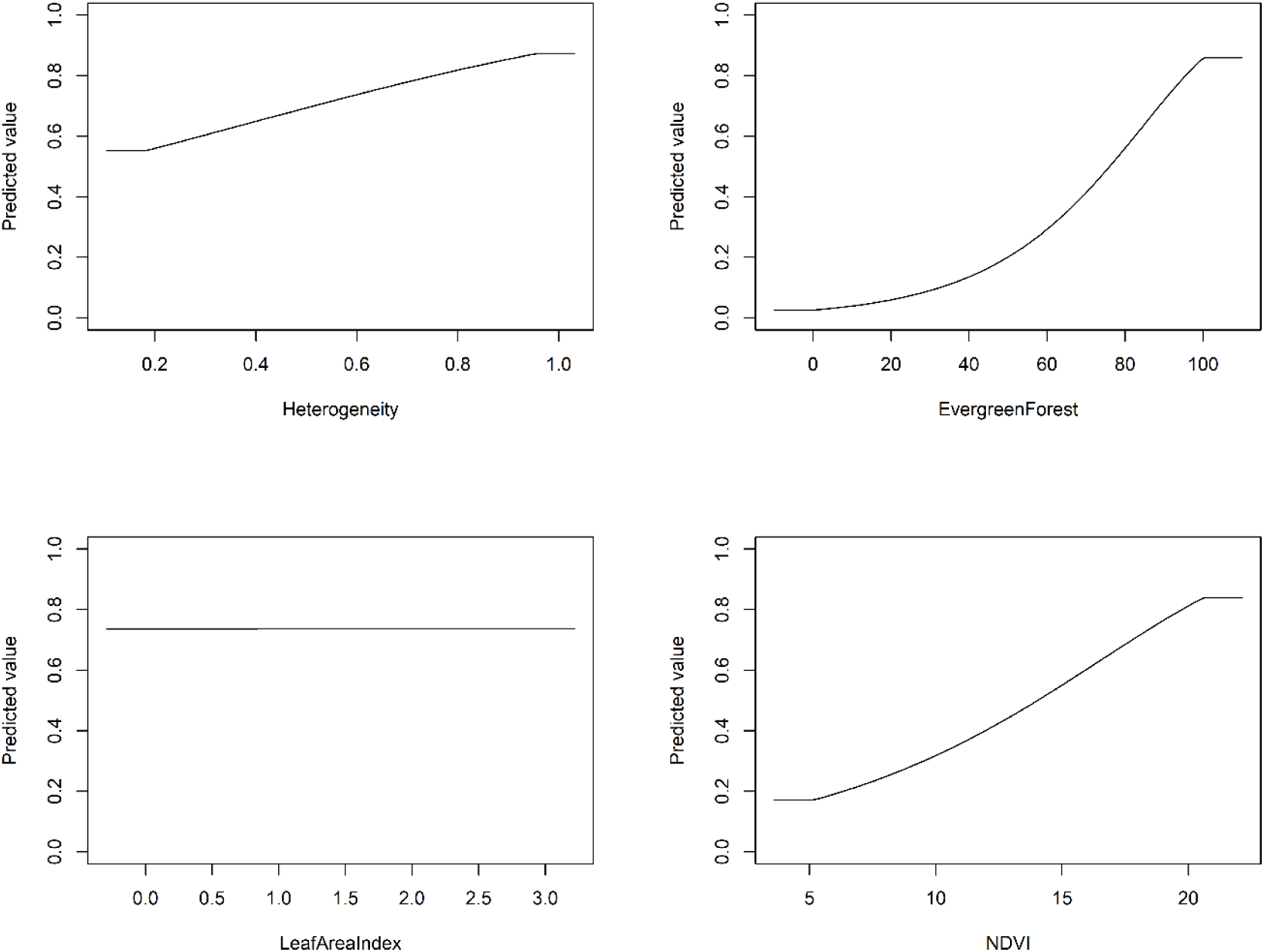
Penalized logistic regression response functions for each habitat covariate from the optimal Species Distribution Model for the Madagascar Serpent-eagle. The curves show the contribution to model prediction (y-axis) as a function of each continuous habitat covariate (x-axis). Maximum values in each response curve define the highest predicted relative suitability. The response curves reflect the partial dependence on predicted suitability for each covariate and the dependencies produced by interactions between the selected covariate and all other covariates.

### Range metrics, population size and protected area coverage

The reclassified binary model (maxTSS threshold = 0.670) calculated a *model* AOH = 30,121 km^2^, 13 % less than the current IUCN range map area of 34,655 km^2^ (Fig. 5). From the *model* AOH, maximum EOO was 397,293 km^2^ and minimum EOO 281,736 km^2^ (Fig. 5), with an AOO = 79,520 km^2^. Using our formulation based on habitat area from nearest neighbour distances, we calculated that the *model* AOH could potentially support 533 mature individuals, or 267 breeding pairs, across the entire Madagascar Serpent-eagle range. The current protected area network covered 95 % (28,654 km^2^) of the *model* AOH, 50 % greater than the target protected area representation of 45 % (Fig. 6). Priority areas of habitat which are without protected area coverage in the protected area network were identified for: (**1**) a large area of forest at Alan’ i Fampanambo linking up to Ambotavoky Special Reserve, (**2**) a forest corridor 20-km west of Anosibe an’ala extending north from Marolambo National Park and (**3**) connecting Midongy Befotaka National Park with d’Andohahela National Park in the far south (Fig. 6, blue circles). 189

**Figure 5.**
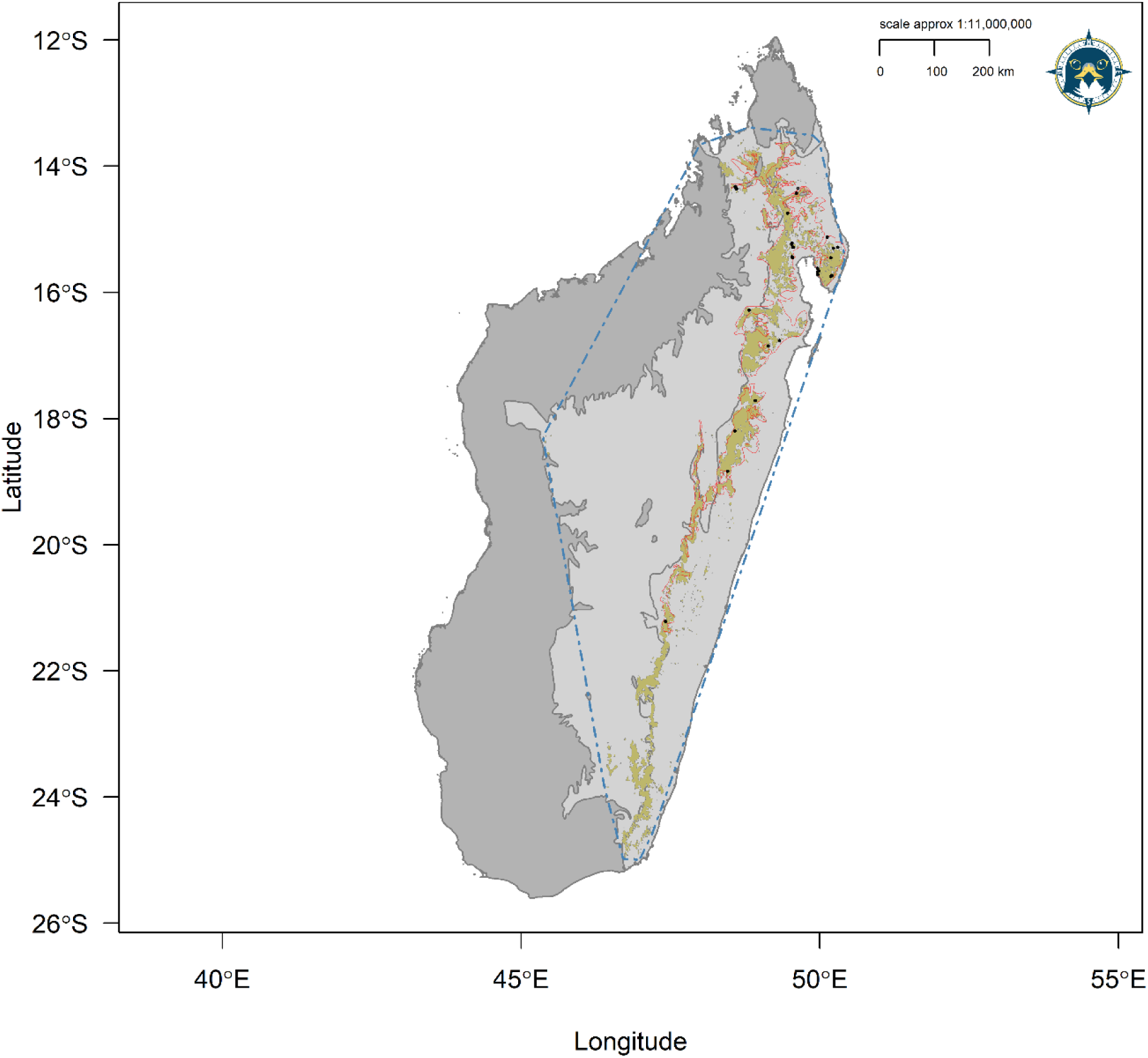
Updated IUCN range metrics for the Madagascar Serpent-eagle showing the reclassified binary *model* area of habitat (AOH, brown polygons) and extent of occurrence (EOO, hashed blue polygon). Red polygon defines current IUCN range map. Light grey polygons represent the species accessible area. Dark grey polygon defines the national boundary of Madagascar not within the species accessible area. Black points define Madagascar Serpent-eagle occurrences from the Global Raptor Impact Network population monitoring system.

**Figure 6.**
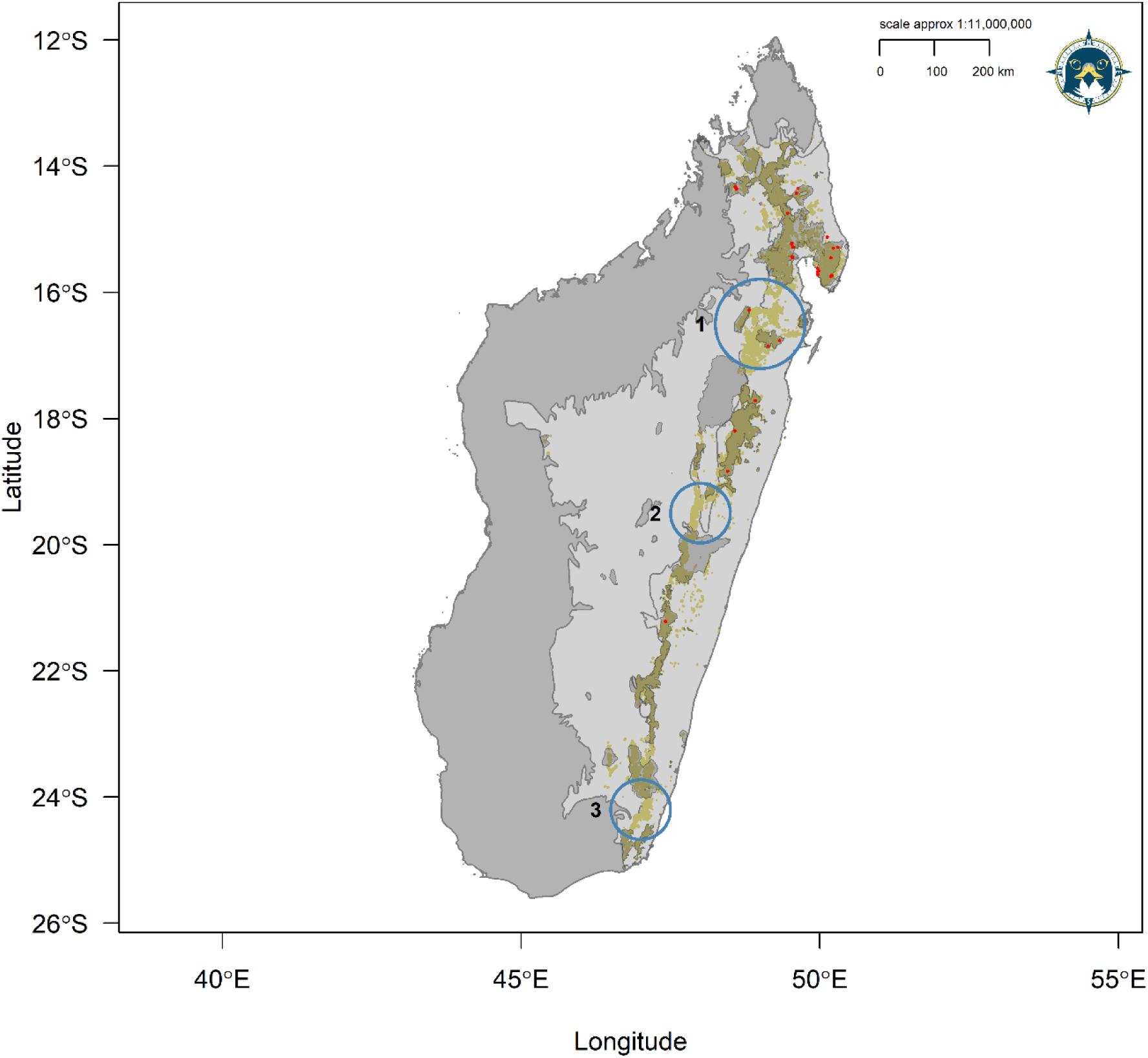
Protected area network coverage for the Madagascar Serpent-eagle showing the reclassified binary *model* area of habitat (AOH, brown polygons) overlaid with the World Database on Protected Areas (WDPA) network (black bordered polygons). Light grey polygons represent the species accessible area. Dark grey polygon defines the national boundary of Madagascar not within the species accessible area. Red points define Madagascar Serpent-eagle occurrences from the Global Raptor Impact Network population monitoring system. Blue circles identify priority WDPA network coverage gaps: (**1**) Alan’ i Fampanambo forest and surrounding area north, (**2**) a forest corridor 20-km west of Anosibe an’ala extending north from Marolambo National Park, and (**3**) a forest corridor connecting Midongy Befotaka National Park with d’Andohahela National Park.

## Discussion

Raptors in developing countries with small geographic ranges that are forest-dependent are particularly extinction-prone and under-studied (Buechley *et al*. 2019). Additionally, tropical forest raptor species are more threatened compared to tropical non-forest raptors, mainly due to habitat alteration driven by logging and land clearance for agriculture (McClure *et al*. 2018). This is further compounded for conservation action by the lack of fundamental biological information on tropical raptors in general (Buechley *et al*. 2019), required for underpinning the scientific understanding required to effect policy and conservation action (McClure *et al*. 2018). The Madagascar Serpent-eagle is thus a prime example of a raptor facing all these combined threats and knowledge gaps. Our results updated previous IUCN range metrics, with our area of habitat map predicting beyond the Madagascar Serpent-eagle known range. We estimated a population size of 533 mature individuals and that 95 % of the area of habitat is currently protected but recommend new protected areas for full protection of habitat across the species range.

Species range metrics are a key component for assessing the conservation status and extinction risk of taxa on the IUCN Red List (IUCN 2019). Using model-based interpolation within our SDM framework we were able to extend the current known range of the Madagascar Serpent-eagle (BirdLife International 2016), predicting into an extensive area further south than the current IUCN range map (See Fig. 5). However, despite this predicted range extension our area of habitat map closely matched that of the IUCN, albeit 13 % less than the IUCN range area. We recommend this updated range map is incorporated into the next Red List assessment for the Madagascar Serpent-eagle. In the meantime, exploratory surveys should be undertaken to assess model accuracy in this newly predicted habitat area, similar to previous SDMs used for rare taxa in Madagascar (Raxworthy *et al*. 2003; Pearson *et al*. 2006).

Quantifying species-habitat associations are key to understanding species’ habitat requirements and environmental preferences (Matthiopoulos *et al*. 2020). We identified the most parsimonious habitat variables based on our occurrence data fitted with multiple logistic regressions. Interestingly, including Climatic Moisture Index resulted in the worst performing habitat covariate model (model #1, Table 2), despite the assumption that climate is key to defining species range limits at continental scales (Pearson & Dawson 2003). We suspect that vegetation indices such as NDVI, which can be strongly correlated with climatic conditions (Ichii *et al*. 2002; Pettorelli 2013), were better able to capture the broad scale tropical forest vegetation dynamics and thus habitat associations for the Madagascar Serpent-eagle. Similarly, topography was not as important when compared to biophysical measures such as Leaf Area Index and FPAR, with neither topographic covariate in the best supported models (Table 2).

From the penalized SDMs, our best model identified the strongest association with vegetation species richness derived from Enhanced Vegetation Index, followed by composite NDVI, concurring with Enhanced Vegetation Index being a more important biophysical measure in dense tropical forests (Huete *et al*. 2002; Qiu *et al*. 2018). Madagascar Serpent-eagles had a flat response up to LAI values of 3.0 (see Fig. 4), concurrent with the negative association in the best fit habitat covariate model. This suggests a weak association between Madagascar Serpent-eagle occurrence with Leaf Area Index values lower than expected from a global analysis (Asner *et al*. 2003), though this study did not include Madagascar. Perhaps the flat to negative association with LAI was related to our low occurrence sample and further compounded by the inclusion of the evergreen forest landcover layer. Importantly, our penalized SDM was able to identify a strong positive association with >95 % evergreen forest cover, following previous ground-based associations (Thorstrom & Rene de Roland 2000; Benjara *et al*. 2021).

Estimating population size is key for IUCN Red List assessments, because it is used in the criteria for designating the specific Red List threat category for a given taxon (IUCN 2019). Our estimate of 533 mature individuals based on predicted AOH is within the population size range currently given by the IUCN (250-999; BirdLife International 2016). Our estimate would technically place the Madagascar Serpent-eagle in the Vulnerable category based on criterion D for a very small or restricted population (IUCN 2019). However, due to low breeding productivity (1 young every 2-3 years) and high juvenile mortality, we are reluctant to recommend re-listing from Endangered to Vulnerable without first assessing population size from further ground-truthing surveys. Encouragingly, protected area coverage was very high, and we recommend consideration of the three major gaps we have identified here as new protected areas, further supported by exploratory surveys to confirm presence. Protected areas have been effective in preventing species extinctions (Geldmann *et al*. 2013). Therefore, protecting as much Madagascar Serpent-eagle habitat as possible is key to its future survival as done previously in the Masoala Peninsula (Thorstrom & Rene de Roland 2000).

We recognise there are limitations to our approach regarding sample size, but we used the current best-practice modelling methodology combined with robust remote sensing variables to calculate our baseline metrics. Even though unstructured occurrence data can have sampling bias (Beck *et al*. 2014), opportunistically collected presence-only data are often the only location data available and generally sample beyond the extent of the smaller spatial scale of systematic surveys (Sutton *et al*. 2020). Thus, when used in conjunction with a modelling framework designed to account for inherent spatial biases unstructured data can fill distributional knowledge gaps (Rhoden *et al*. 2017; Sutton *et al*. 2022). However, obtaining further occurrences would be useful for improving our predictions and updating the baseline biological parameters set out here.

Madagascar has been identified as a priority region for raptor research and conservation, due to its range of endemic, under-studied raptors (Buechley *et al*. 2019). Future modelling goals include predicting the core remaining areas of habitat for all Madagascar raptors to identify priority areas for current spatial conservation planning. Future work should thus focus on building upon the SDM framework set out here to estimate range metrics, population size, and protected area coverage for all Madagascar raptors combined with remote sensing technology. Our model framework is a fast, cost-effective method to establish key spatial conservation baselines. This framework is widely applicable across all taxa but particularly for rare, under-studied species such as the Madagascar Serpent-eagle which face threats to their future survival. 1047 5018

## Acknowledgements

We thank all individuals and organisations who contributed occurrence data to the Global Raptor Impact Network (GRIN) information system. We thank the M.J. Murdock Charitable Trust for funding and technicians from The Peregrine Fund’s Madagascar Project for support and fieldwork: Ladoany Eugene, Moïse and Monesy

## Data Accessibility Statement

Upon acceptance the data that support this study will be made openly available on the data repository *figshare*

## Authors’ Contribution Statement

*LJS* conceived the idea and designed methodology; *AB, LARR and RT* collected the data; *LJS* analysed the data and led the writing of the manuscript with supervision from *CJWM*. All authors contributed critically to the drafts and gave final approval for publication.

## Supplementary Material

### Methods

We cleaned community science occurrences by removing duplicate records, those with no georeferenced location, and any locations over water, resulting in the 33 occurrences to use in the calibration models (Fig. 1). Further, community science data were checked for geolocation accuracy and any locations that were deemed biologically implausible due to being outside the species’ known habitat requirements were removed from the analysis. Only locations recorded from 1990 onwards were used to match the timeframe of the environmental raster data, whilst retaining sufficient sample size for modelling (van Proosdij *et al*. 2016). We omitted occurrences originating from the eBird database (*n =* 5, Sullivan *et al*. 2009) from the GBIF download because of uncertainty in location accuracy from the eBird checklist system.

We tested spatial clustering in our point locations using Nearest Neighbour Index (NNI) to evaluate the potential level of sampling bias. NNI is the ratio of the observed distance divided by the expected distance between neighbours in a hypothetical random distribution. NNI < 1 indicates spatial clustering, with values of 1 indicating random dispersion, and those closer to 2 indicating regular dispersion. Despite the advantages gained from applying spatial filtering to occurrence data in SDMs (Aiello-Lammens *et al*. 2015), we opted to retain all 38 cleaned records without spatial filtering because the effects of biased sampling were minimal in our dataset (NNI = 0.879, *z* = −1.33, *p =* 0.182). By retaining all records we aimed to capture the widest possible range of species-habitat associations because of the low sample size of our occurrence data and the high environmental heterogeneity of the study area.

Continuous Boyce Index (CBI) is consistent with a Spearman correlation (*r_s_*) with CBI values ranging from −1 to +1, with positive values indicating predictions consistent with observed presences, values close to zero no different than a random model, and negative values indicating areas with frequent presences having low environmental suitability. CBI was calculated using 20 % test data with a moving wi ndow for threshold-independence and 101 defined bins in the R package enmSdm (Smith 2019). We Partial ROC ratios range from 0 – 2 with 1 indicating a random model. Function parameters were set with a 10 % omission error rate, and 1000 bootstrap replicates on 50 % test data to determine significant (𝛼 = 0.05) pROC values >1.0 in the R package ENMGadgets (Barve & Barve, 2013).

## Supplementary Tables

**Table S1.**
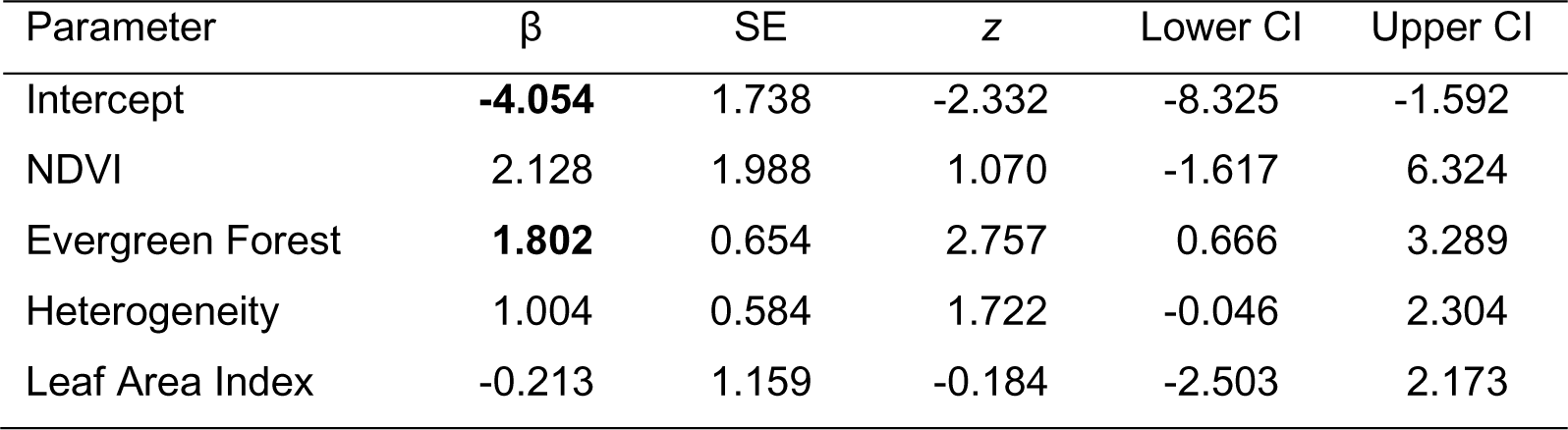
Model parameters from the best supported candidate logistic regression habitat covariate model (#6) for the Madagascar Serpent-eagle using Akaike’s Information Criterion corrected for small sample sizes. Bold indicates significant coefficient estimates *p* < 0.01.

**Table S2.** Multi-collinearity test using stepwise elimination Variance Inflation Factor (VIF) analysis. Covariates with VIF < 2 have low correlation with other covariates, and thus are suitable for inclusion in calibration models when further evaluated for ecological relevance.

| Covariate | $R^2$ | Tolerance | VIF |
| --- | --- | --- | --- |
| Heterogeneity | 0.476 | 0.524 | 1.909 |
| NDVI | 0.399 | 0.601 | 1.664 |
| Evergreen Forest | 0.274 | 0.726 | 1.377 |
| Leaf Area Index | 0.257 | 0.743 | 1.347 |

**Table S3.**
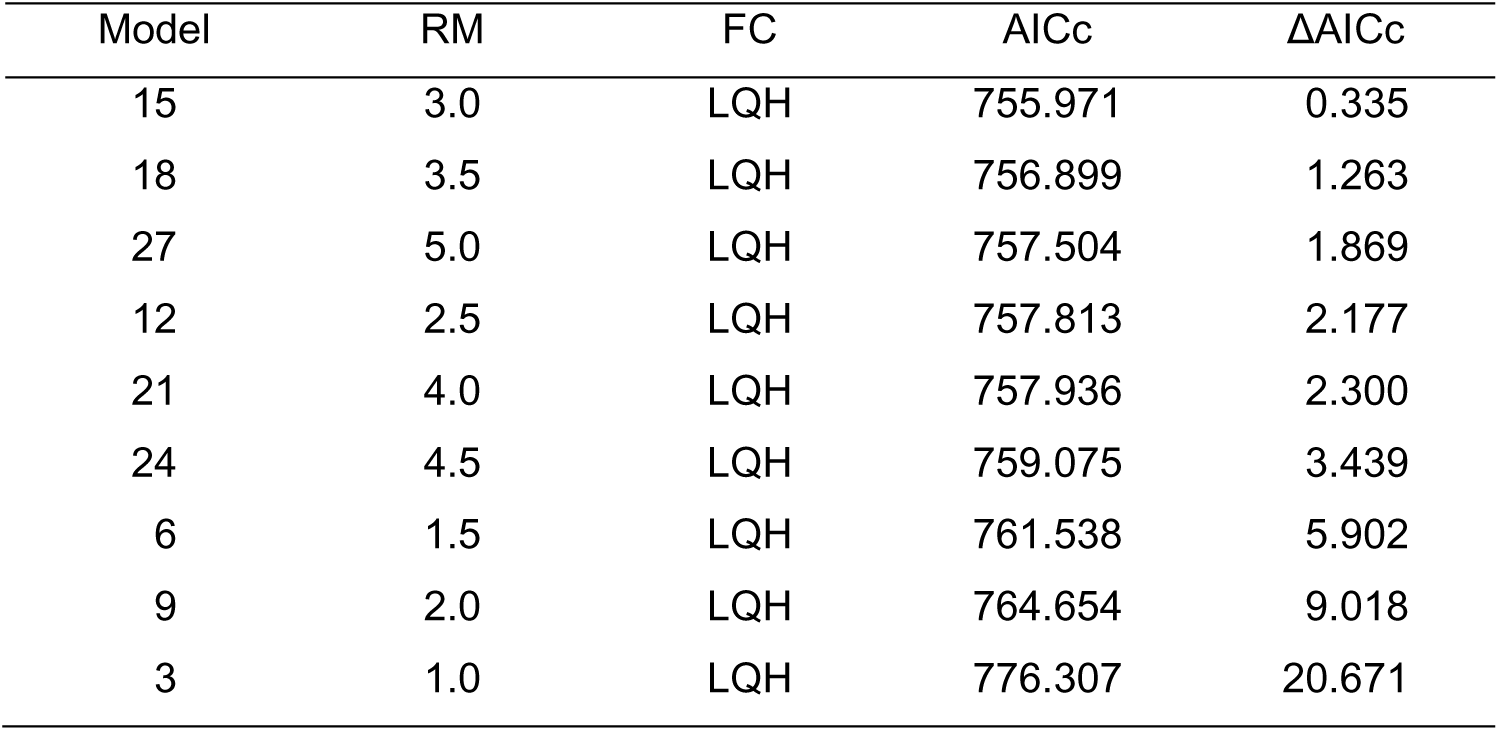
Model selection metrics for all candidate models using all three feature classes from Akaike’s Information Criterion corrected for small sample sizes. RM = regularization multiplier (β), FC = feature classes, LQH = Linear, Quadratic, Hinge.

## Supplementary Figures

**Figure S1.**
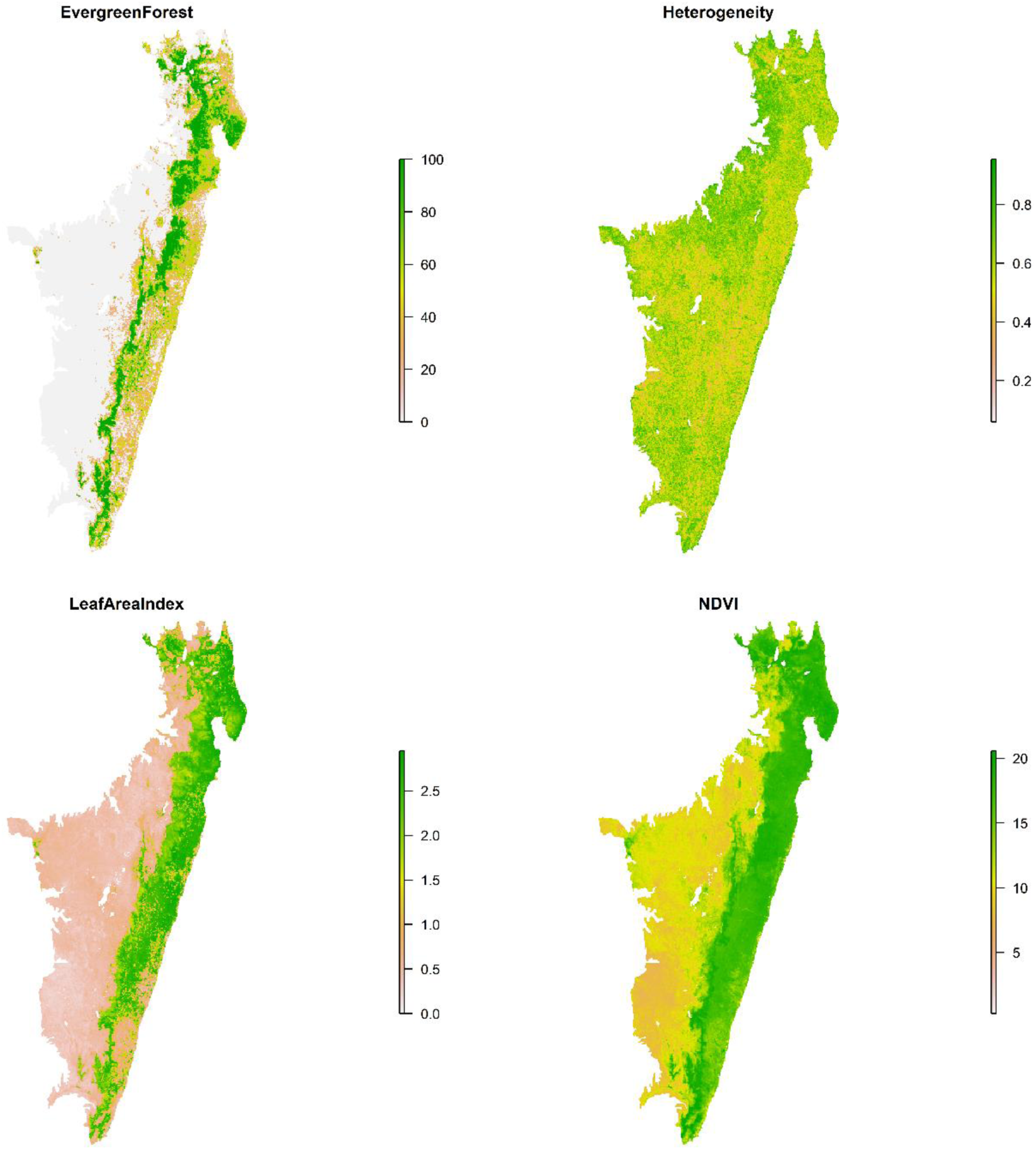
Habitat covariates used in Species Distribution Models for the Madagascar Serpent-eagle.

**Figure S2.**
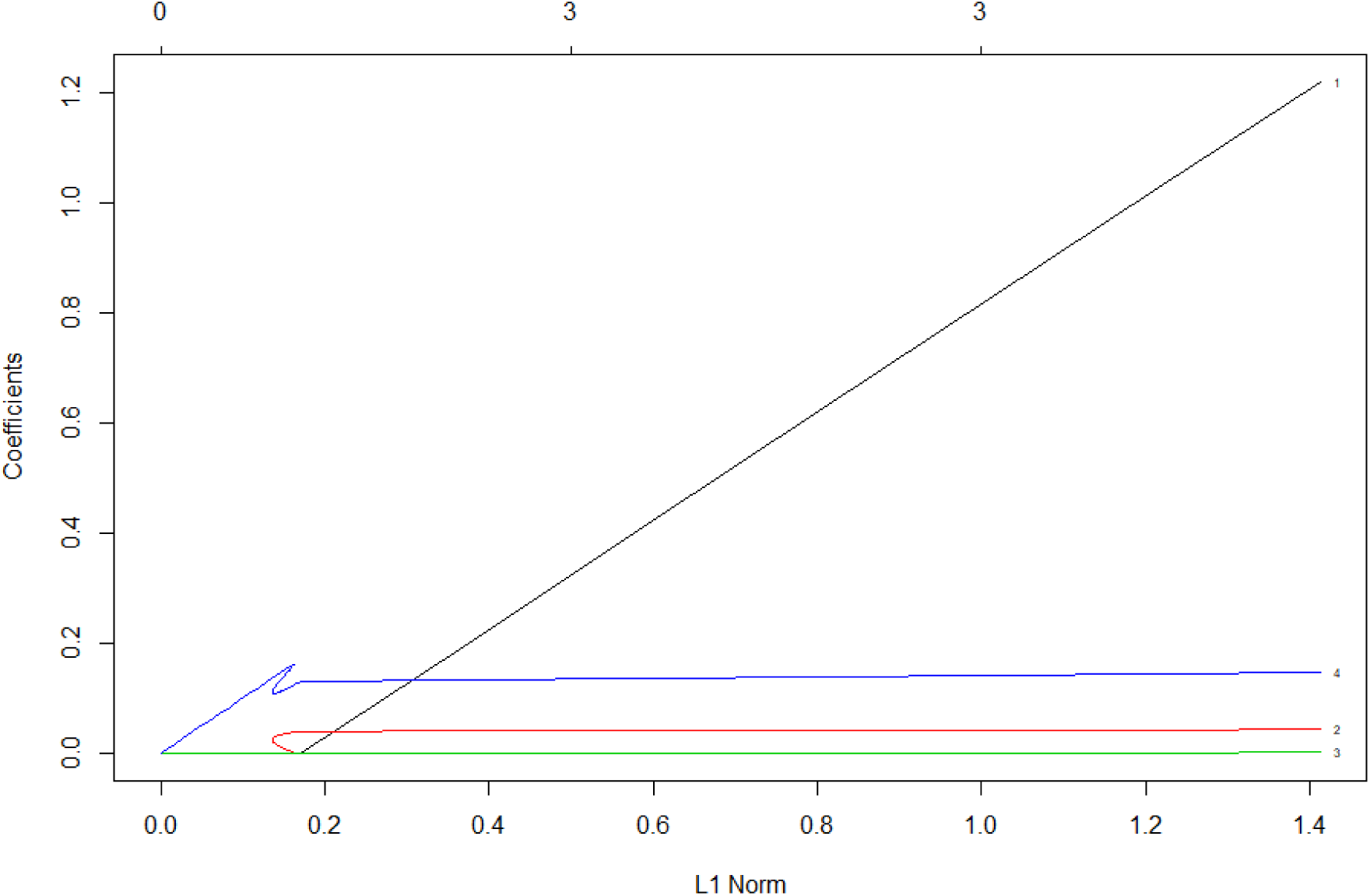
Beta coefficient paths for the optimal penalized logistic regression model where each curve corresponds to a covariate linear term. The paths of each coefficient term are plotted against the L1-norm (lasso or elastic net) of the whole coefficient vector as lambda (the amount defining the level of coefficient shrinkage) varies. The upper axis indicates the number of non-zero coefficients at the current lambda which is the effective degrees of freedom for the lasso or elastic net.

**Figure S3.**
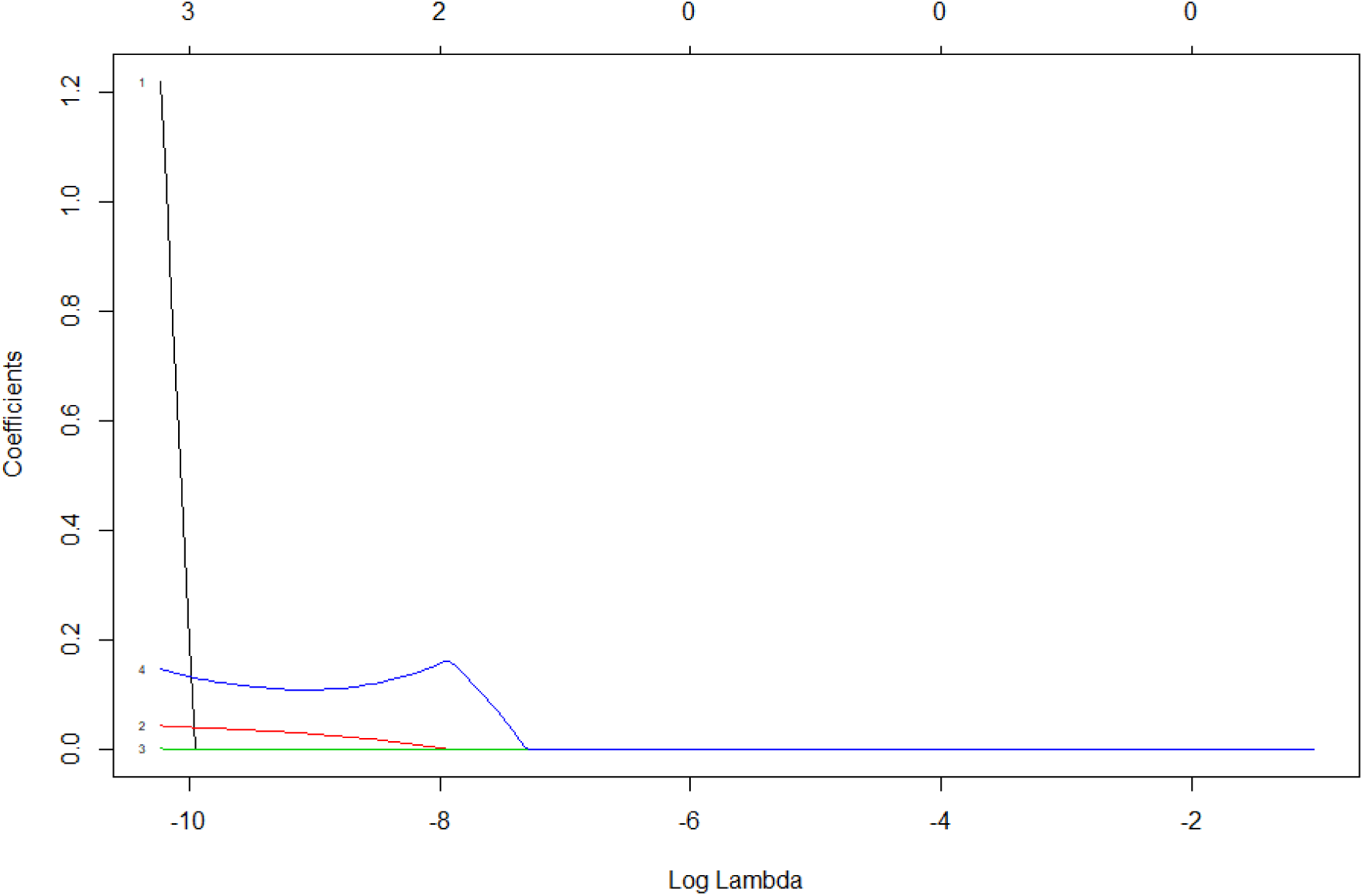
Beta coefficient paths for the optimal penalized logistic regression model where each curve corresponds to a covariate linear term. The paths of each coefficient term are plotted against the log-lambda of the whole coefficient vector as lambda (the amount defining the level of coefficient shrinkage) varies. Log-lambda on the y-axis indicates the log of the optimal value of lambda which minimizes the prediction error. This lambda value will give the most accurate model. The upper axis indicates the number of decreasing non-zero coefficients at the current lambda which is the effective degrees of freedom for the elastic net.

**Figure S4.**
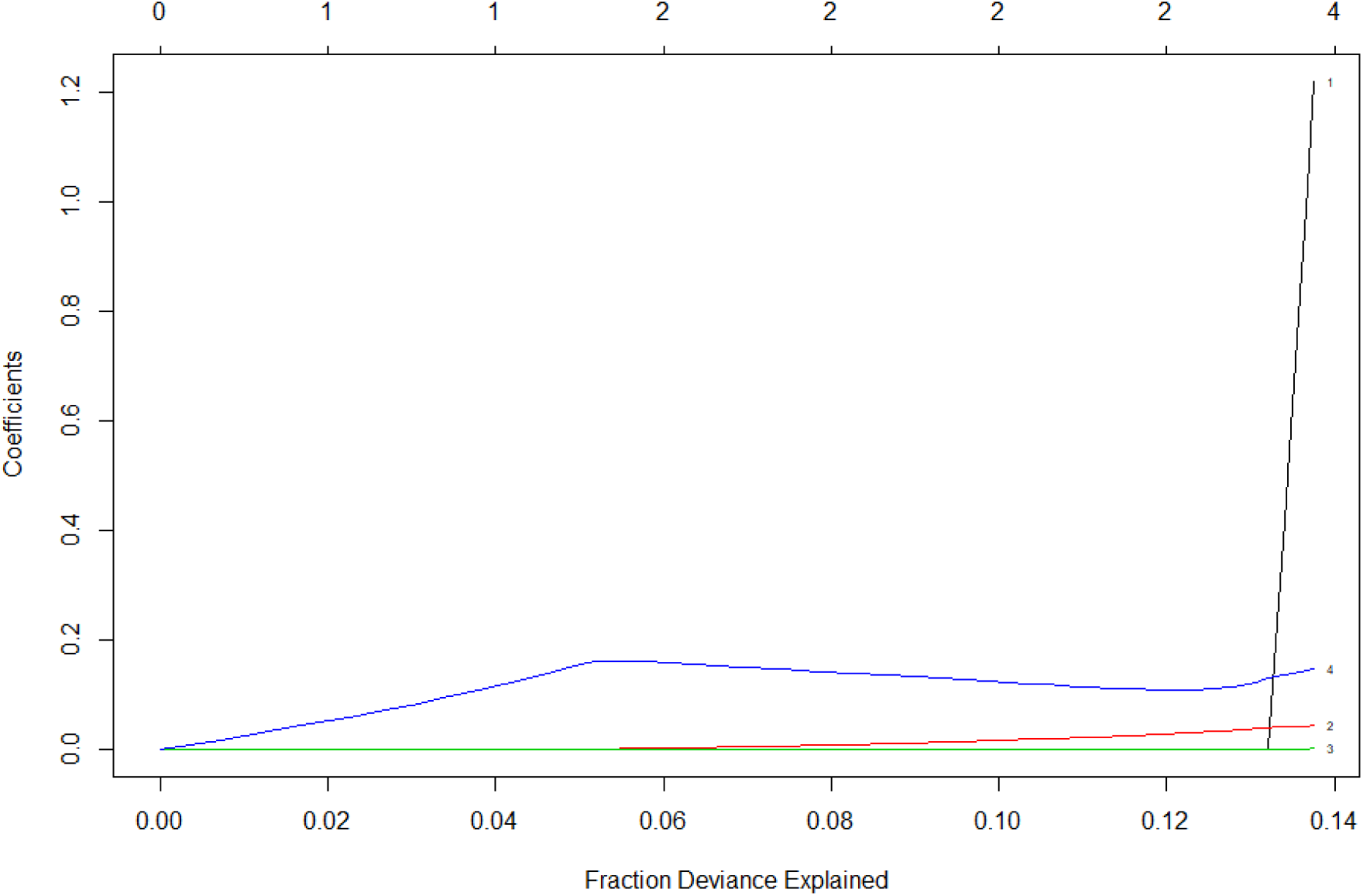
Beta coefficient paths for the optimal penalized logistic regression model where each curve corresponds to a covariate linear term The paths of each coefficient term are plotted against the fraction deviance explained on the training data. The upper axis indicates the number of non-zero coefficients at the current lambda which is the effective degrees of freedom for the elastic net.

